# Real-time assembly of an artificial virus elucidated at the single-particle level

**DOI:** 10.1101/526046

**Authors:** Margherita Marchetti, Douwe Kamsma, Ernesto Cazares Vargas, Armando Hernandez García, Paul van der Schoot, Renko de Vries, Gijs J.L. Wuite, Wouter H. Roos

## Abstract

While the structure of a variety of viruses has been resolved at atomistic detail, their assembly pathways remain largely elusive. Key unresolved issues in assembly are the nature of the critical nucleus starting particle growth, the subsequent self-assembly reaction and the manner in which the viral genome is compacted. These issues are difficult to address in bulk approaches and are effectively only accessible by tracking the dynamics of assembly of individual particles in real time, as we show here. With a combination of single-molecule techniques we study the assembly into rod-shaped virus-like particles (VLPs) of artificial capsid polypeptides, de-novo designed previously. Using fluorescence optical tweezers we establish that oligomers that have pre-assembled in solution bind to our DNA template. If the oligomer is smaller than a pentamer, it performs one-dimensional diffusion along the DNA, but pentamers and larger oligomers are essentially immobile and nucleate VLP growth. Next, using real-time multiplexed acoustic force spectroscopy, we show that DNA is compacted in regular steps during VLP growth. These steps, of ∼30 nm of DNA contour length, fit with a DNA packaging mechanism based on helical wrapping of the DNA around the central protein core of the VLP. By revealing how real-time, single particle tracking of VLP assembly lays bare nucleation and growth principles, our work opens the doors to a new fundamental understanding of the complex assembly pathways of natural virus particles.

## Introduction

The structure of a large class of viruses is highly regular and stable (1), and a number of these regular viruses have been reconstituted *in vitro*, suggesting that physical driving forces determine the assembly process. This motivated trials in which the viral genome is replaced by other cargo molecules, allowing viruses to be employed, e.g., as therapeutic platforms for the delivery of polynucleotides and drugs (2–4). Because of their importance in health care, nanomedicine and nanotechnology, a variety of bulk experimental studies (5–8) as well as modelling and computer simulation approaches (9–11) have been performed to attempt to elucidate the mechanism underlying viral assembly. This has yielded insights on possible assembly mechanisms albeit that the multiplicity of assembly pathways in the experiments obscure interpretation of the findings (12). Key issues yet to be resolved are the nature of the critical nuclei required for productive viral capsid formation, as well as the nature and dynamics of nucleic acid condensation during capsid formation (13–15). In order to discriminate between different pathways and to identify assembly intermediates, single particle techniques are needed.

Single molecule techniques such as electron microscopy and atomic force microscopy (AFM) provide for high resolution images of viruses, but typically yield images with only limited information on viral dynamics (16, 17). High resolution AFM imaging and electron microscopy have recently shown transient capsid intermediates, from which assembly kinetic parameters can be estimated (18). New approaches such as resistive-pulse sensing, optical tweezers and acoustic force spectroscopy have been reported to study the assembly of the icosahedral viruses Hepatitis B Virus and SV40 virus (19, 20). Here we go beyond these recent studies by using combined confocal fluorescence and optical tweezing to identify the nature of critical nuclei in capsid formation, and acoustic force spectroscopy, to probe the dynamics and nature of nucleic acid condensation during the formation of single, rod-shaped capsid particles in real time. We reveal the assembly kinetics of a previously de-novo designed artificial capsid polypeptide (21), which co-assembles with single and double-stranded DNA templates into virus-like particles (VLPs). These rod-shaped VLPs consist of a single DNA molecule coated with multiple copies of the artificial capsid polypeptide. The sequence of the artificial capsid polypeptide was designed to mimic the co-assembly mechanism of the Tobacco Mosaic Virus (TMV) capsid protein with its nucleic acid template (21–23). The polypeptide, referred to as C-S_10_-B, consists of three blocks that each encode a specific physico-chemical functionality, mimicking corresponding functionalities of viral capsid proteins **(Fig. 1A)**.

**Fig 1.**
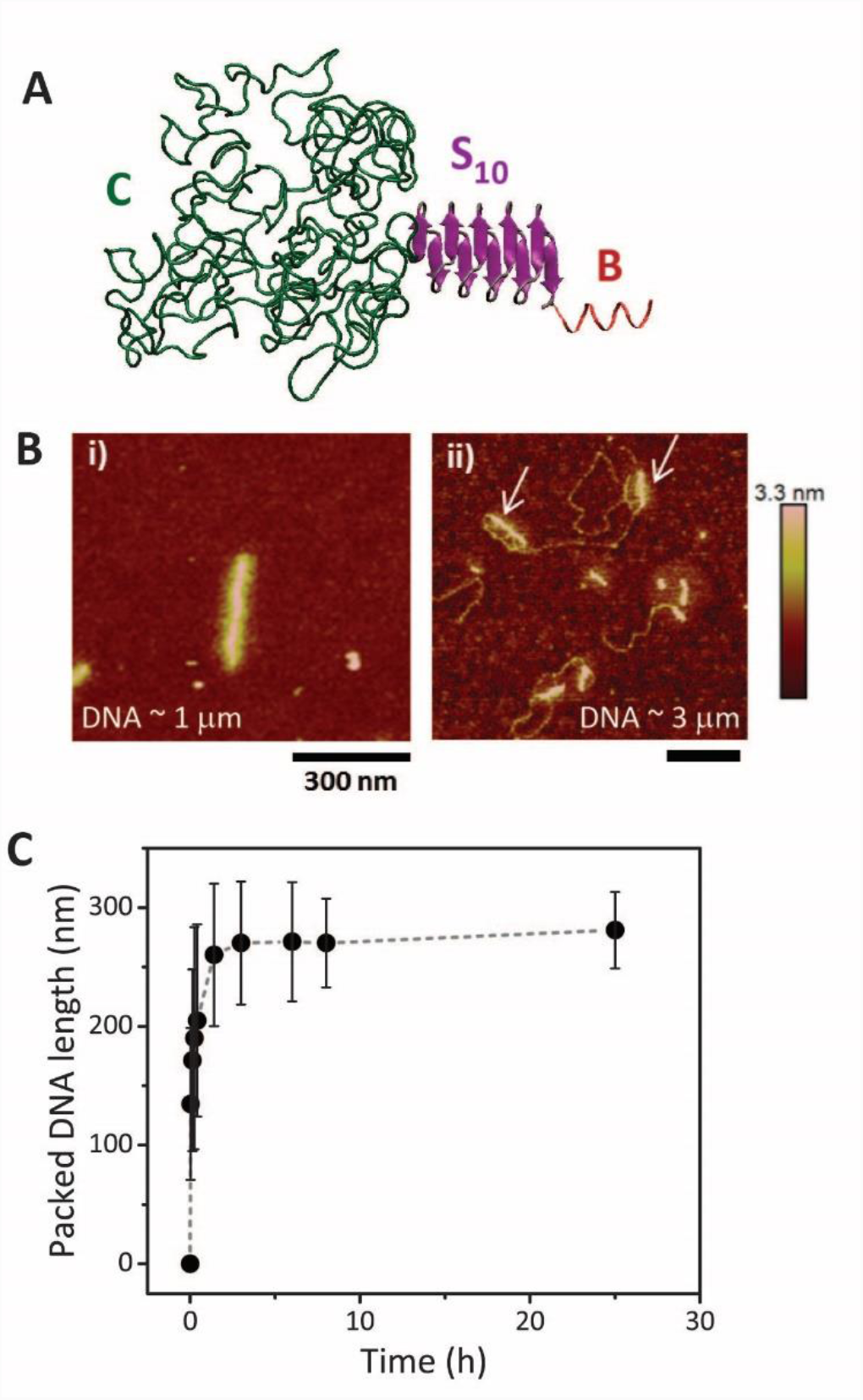
The artificial capsid polypeptide under investigation and the resulting formation of rod-shaped particles. **(A)** A schematic of the tri-blocks polypeptide C-S_10-_B. Each block is highlighted by a different color and its specific function, related to its physiochemical properties, is described in the main text. **(B)** Particles formation on DNA molecules of different lengths was probed with AFM imaging in air. On DNA of ≈1 μm contour length mainly single particles form in which the DNA is compacted 1/3 of its original length (panel (i)) (see also SI). When a 3-folds longer DNA is employed (∽3 μm contour length) 2-4 nucleation points are observed in the early stages of particles assembly (panel (ii)). The white arrows point at two different nucleation sites that are formed on the same DNA molecule. (C) Quantification from AFM images of the ‘slow’ kinetics formation of particles on a 2.5 kbp DNA, being packed to 1/3 of its original length, as also previously shown (21).

Nucleic acid binding is achieved through interactions with block B that consists of 12 positively charged lysines. The silk-like middle blocks S_10_ = (GAGAGAGQ)_10_ fold into a sheet-like beta-solenoid conformation (23, 24) **(Fig. 1A)**, and stacking of these sheets leads to the formation of a rigid protein filament that forms the core of the VLP (21). Folding of an initially unfolded silk block into the beta-solenoid conformation is promoted by docking onto an already existing folded silk block, such that the formation of the rod-shaped protein core is a nucleated process (20, 21, 35). Finally, a hydrophilic random-coil C, with a collagen-like sequence C = (GXY)_132_ (where X and Y are mostly hydrophilic uncharged amino acids (26)) provides colloidal stability to the rod-shaped VLPs. Immediately after dissolution, the silk blocks of C-S_10_-B polypeptides are still unfolded, but over time they fold and stack, a process that is strongly promoted by binding to the nucleic acid templates, such that co-assembly with nucleic acid templates is favored over capsid protein-only assembly (21).

In bulk, the nucleation and growth of the VLP particles obeys the same simple kinetic model that was earlier shown to describe the nucleation and growth of TMV particles (9, 21, 22). These artificial capsid polypeptides, which can be easily modified, are therefore an ideal model system for studying viral assembly pathways. We here use them to address key issues regarding capsid assembly pathways that are difficult to catch other than with real-time, single particle methods: the nature of the critical nuclei for productive capsid formation, and the dynamics and nature of nucleic acid condensation during capsid formation. Our study on this simple model system paves the way for the detailed real-time *in vitro* studies of the assembly of natural viruses at the single-particle level.

## Results

### Real-time observation of nucleation of multiple artificial capsids on long DNA

First we use Atomic Force Microscope (AFM) imaging to recapitulate basic properties of encapsulation of linear DNA by the artificial capsid polypeptides (17). Upon mixing the capsid polypeptides with double-stranded DNA, rod-shaped particles are formed, confirming earlier findings (21) (**Fig. 1B**). The kinetics of particle formation can be quantified by analysing the length of packaged DNA as a function of time (**Fig. 1C**). DNA with a contour length < 1 μm typically displays one or two nucleation sites whilst longer DNA with a contour length > 1 μm often shows more than two nucleation sites (**Fig. 1B and SI**). The relative frequency of the number of nucleation points and the resulting branches in the self-assembled particle were quantified for a 2.5 kbp-long DNA (contour length of ≈ 850 nm) (**Fig. S1**). In order to estimate the solution diameter of the VLPs, we additionally performed AFM imaging in liquid, finding an average diameter of the VLPs of 9 nm, a value that matches the expected dimensions of the VLP (**Fig. S1**).

Next we turn to studying assembly of single virus-like particles in real-time, first considering the nucleation of artificial capsids on their DNA templates. Specifically, we wish to elucidate the nature of the critical nuclei required for productive capsid growth. In order to do so, we use combined confocal fluorescence microscopy and optical tweezers (27). A long DNA molecule (λ-phage DNA, contour length ≈16.5 μm) is attached at both ends to a microsphere (“bead’’), and both beads are trapped using a double optical tweezers set-up. Simultaneous confocal scanning laser microscopy allows for real-time probing of the local binding of fluorescently labeled artificial capsid polypeptides on the DNA template (28) **(Fig. 2A)**. Because the used DNA in this assay is long multiple nuclei form (**Fig. S1**), making this technique is particularly well suited to zoom in on nucleation events of the VLPs.

**Fig 2.**
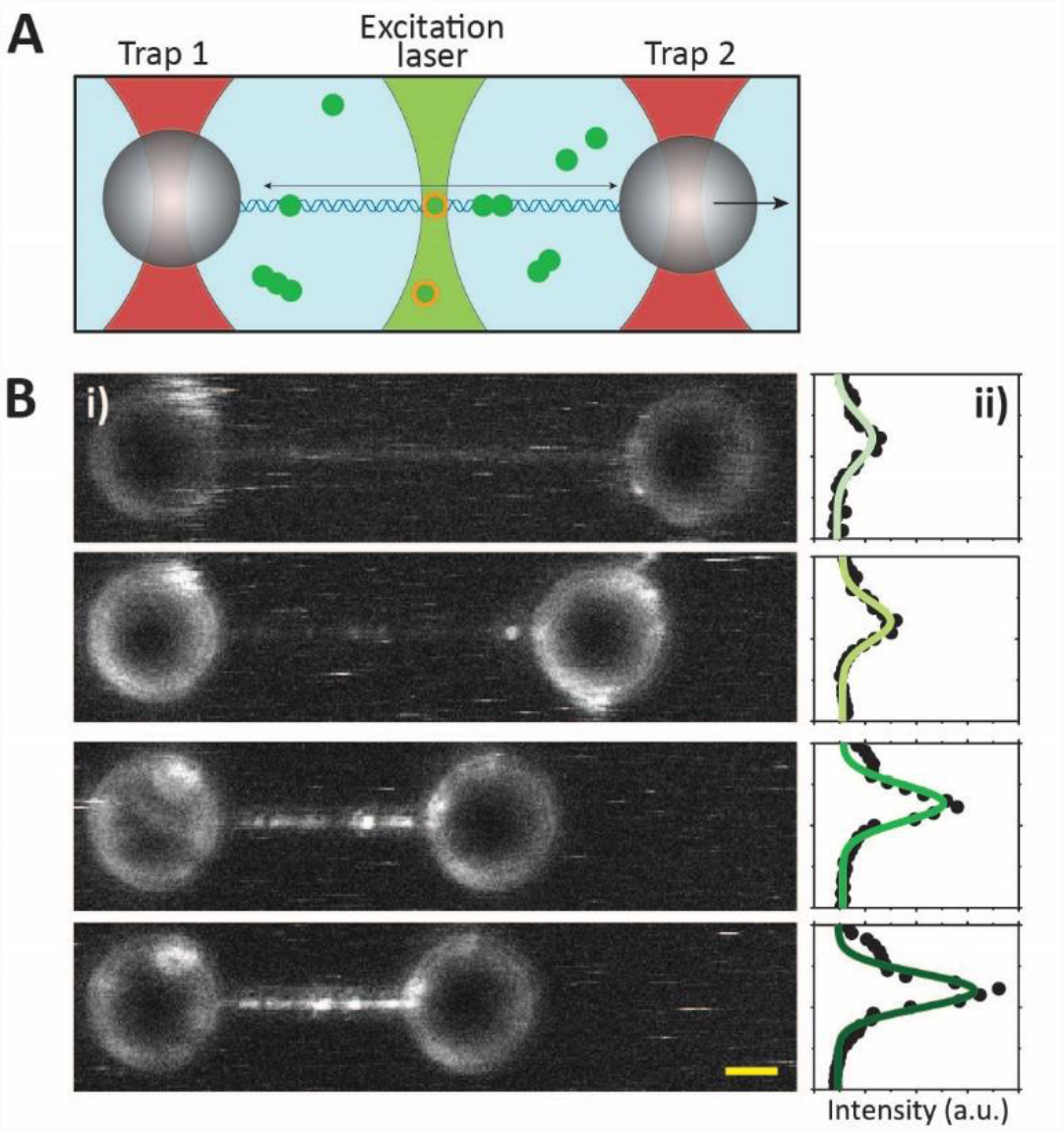
Optical tweezers combined with confocal fluorescence microscopy, protein binding and compaction visualization. **(A)** Illustrative image of a dual trap optical tweezers combined with confocal fluoresce microscopy. Two focused lasers beams (red beams) trap two microspheres that are chemically attached to a DNA molecule. Proteins in solution (green dots) bind to the DNA and their fluorescence tag lights up when the scanning laser (light green beam) illuminates them. **(B)** Progressive packaging of the DNA by the polypeptides. Fluorescence images show how DNA shortening (confocal images, left panel) is accompanied by an increase in the fluorescence intensity given by increasing number of bound peptides (plots in right panel).

First, nucleation is allowed to proceed unimpeded by repeatedly keeping the DNA in a relaxed state (< 1 pN) for a fixed time (5 min), followed by a short period of imaging at a constant force of 5 pN. (**Fig. 2B**). We observe a shortening of the end-to-end distance as a function of time, indicating condensation of the DNA during the relaxed state phases **(Fig. 2B, i)**. We also observe an increasing number of nuclei as a function of time, and a corresponding increase of the total fluorescence intensity **(Fig. 2B, ii)**. With these findings, which are supported by the AFM data, we confirm that in our dynamic assay indeed multiple nucleation sites are formed, capable of compacting the DNA in a progressive way. Next, we stretch the DNA to study the mechanical effect of polypeptide binding to the DNA. For this purpose, the relaxed DNA was first incubated with the proteins, then stretched to an end-to-end distance nearly equal to its contour length, and subsequently relaxed back to zero force while recording the retraction force (**Fig. 3A**). We repeat this relaxation and pulling process on the same DNA molecule, until the DNA is saturated and no additional polypetides are bound. During extension several drops in the force can be observed, indicating that compaction is partially being reversed for increasing pulling force. Furthermore, as the compaction progresses, the retraction force becomes much larger than that for bare DNA. Moreover, when performing two consecutive force-distance curves without incubation time in between, we find that the DNA is again compacted (**Fig. S2**). This indicates that the polypeptides remain associated with the DNA during stretching (see also kymograph in **Fig. S2**), and that they are able to re-compact the DNA once the genome is again in the relaxed state.

**Fig 3.**
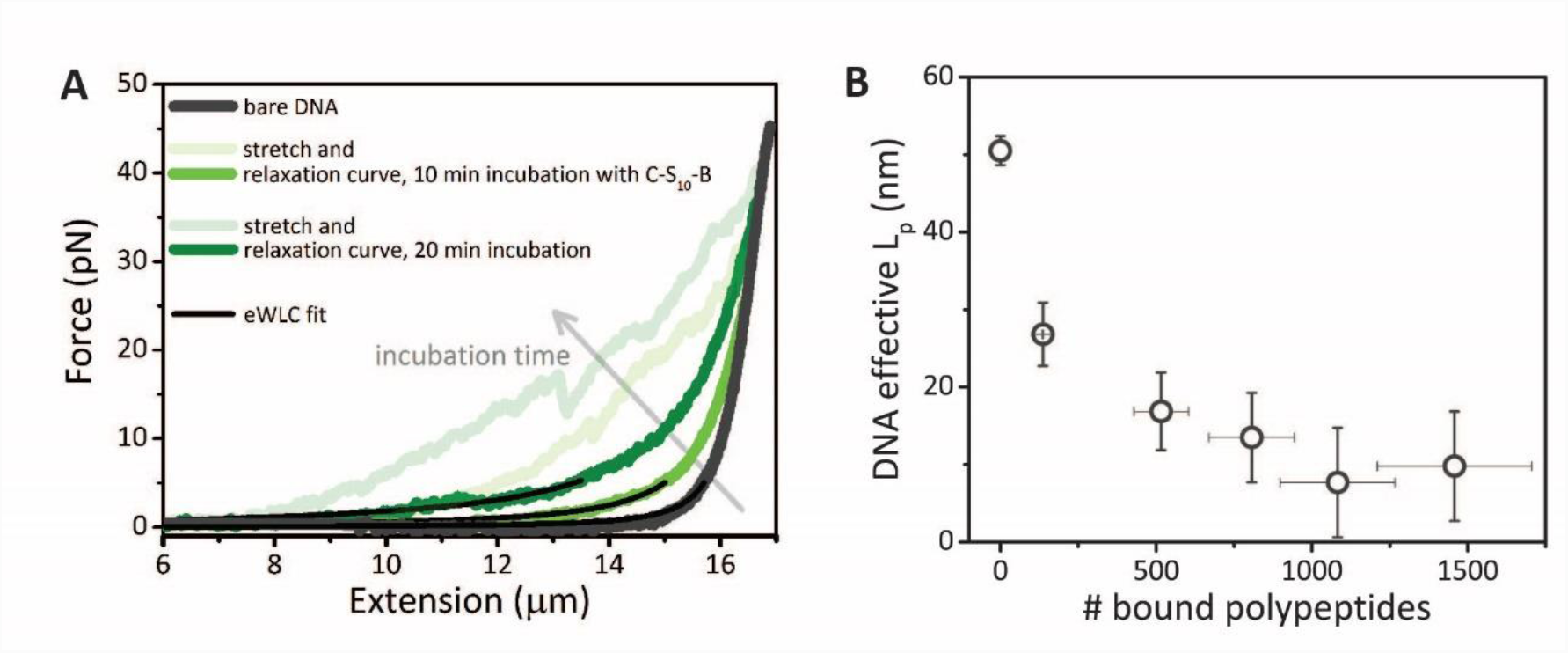
Stretching allows to probe the DNA mechanical changes. **(A)** Repetitive stretching and relaxation of the same DNA molecule in the presence of polypeptides. Representative force-distance curves (FDCs) after different incubation times with 200 nM C-S_10-_B. As a control extension and retraction of bare DNA is shown in dark grey, as expected both curves fall on top of each other. **(B)** The DNA effective persistence length (L_p_) is obtained by fitting the WLC model to the retraction curves. The effective L_p_ decreases as a function of increasing numbers of bound polypeptides. Data point are averages of 15 FD curves at 200 nM of C-S_10-_ B in solution. Error bars correspond to SEM. The errors on x-axis relate to the uncertainty of the estimated intensity of a single fluorophore.

Fitting a worm-like chain model to the retraction curves yields an effective persistence length (L_p_) of the DNA-polypeptide complex (29). In addition, we calibrated the fluorescence intensity of one dye (11.1 ± 0.5 photons, see Methods), in order to determine the number of polypeptides bound. We find that L_p_ decreases sharply as the number of bound capsid polypeptides increases (**Fig. 3B**). Indeed, induced deformations of DNA by polypeptide binding, such as kinking or bending typically give rise to increased retraction forces (30, 31), which can be interpreted as an enhanced apparent flexibility due to the induced (non-thermal) DNA deformations. We have also ascertained that DNA deformations are not caused by the oligolysine binding blocks B *per se*, but require the formation of the rigid protein core of the VLP by the silk-like midblock (S_10_). When performing the force-extension measurements with a polypeptide C-B lacking the central S_10_ silk-like block that is known to simply coat but not condense the DNA (32), there is no DNA shortening and the effective persistence length of the DNA remains unaffected (**Fig. S2**).

### Identification of the nature of critical nuclei required for artificial capsid formation

In the experiments discussed above, nucleation and growth is allowed to proceed unimpeded as the DNA is in a relaxed state for fixed times, and is only stretched for a short time to allow for imaging. This precludes the observation of capsid nucleation with high temporal resolution. Therefore, we next performed experiments to quantify polypeptide binding dynamics at the single molecule level and at millisecond time scales. We keep the DNA at a fixed end-to-end distance of 15.5 μm and continuously monitor the fluorescence along the 16.5 μm contour length long DNA in the form of kymographs **(Fig. 4A)**. In this situation capsid formation is likely to be limited on the stretched DNA while binding dynamics should be unaffected. The calibration of the fluorescence intensity per dye molecule allows us to determine the number of polypeptides involved in each binding event recorded in the kymographs. In **figure 4B** the cumulative binding of capsid polypeptides is plotted for two different polypeptide concentrations. We have used a simple reversible Langmuir adsorption kinetics model (33) to describe peptide adsorption onto the DNA (**SI**). There is a fair agreement for the two different protein concentrations resulting in an equilibrium binding constant K ∼ 7·10^8^ M^−1^. This value corresponds to an effective binding free energy of ∼25 times the thermal energy, i.e., sufficiently strong to withstand typical thermally induced rupture. Furthermore, we find the adsorption rates to scale linearly with concentration, suggesting a diffusion-limited binding process. The strong electrostatic binding is also confirmed by stretching to high forces of polypeptide-coated DNA, where no unbinding is observed (**Fig. S2**).

**Fig 4.**
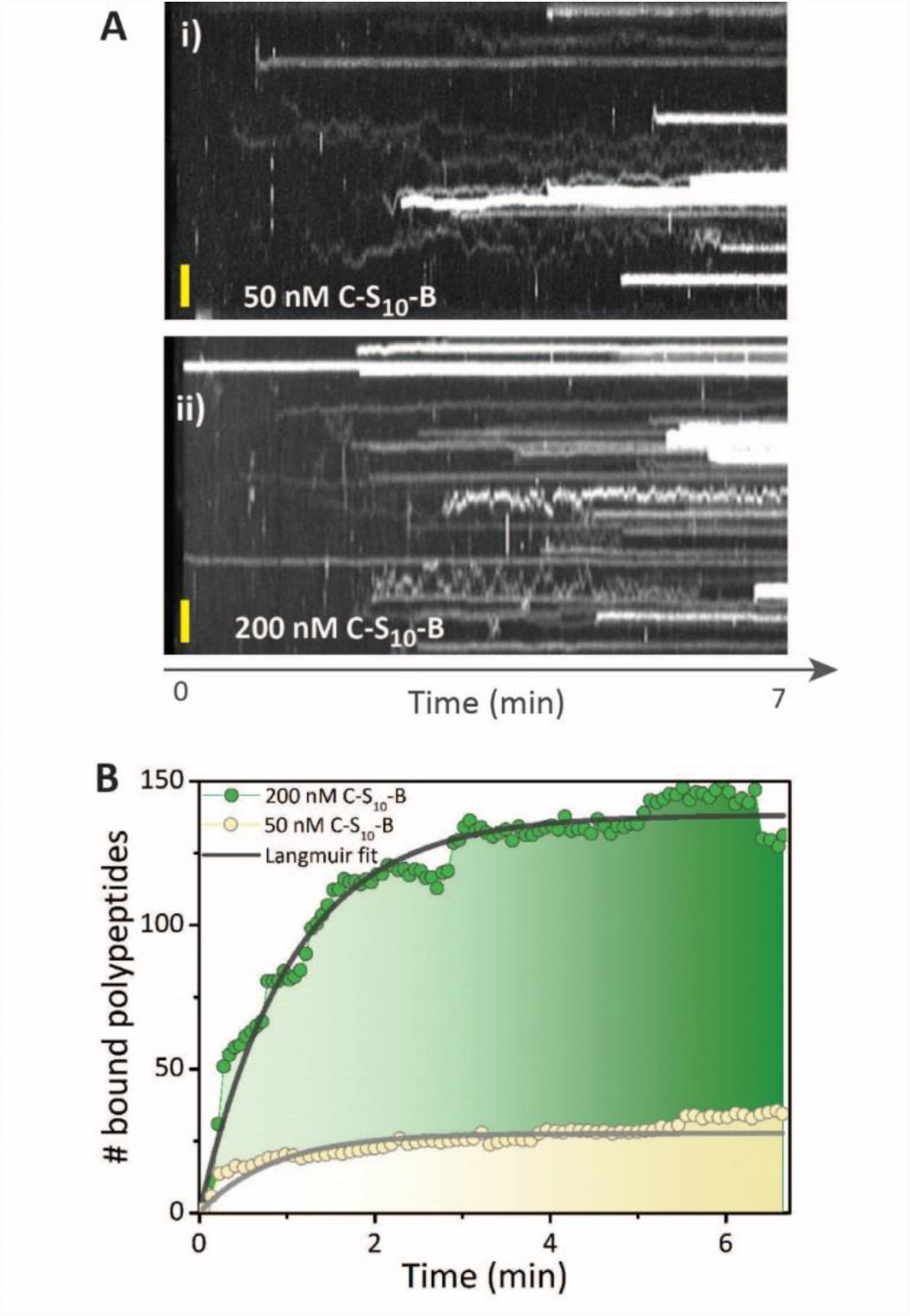
Real-time polypeptide binding. **(A)** Kymographs showing progressive peptide binding for two different C-S_10-_B concentrations: 50 nM panel (i) and 200 nM panel (ii). The confocal scanning line-time is 30 ms, the yellow scale bar denotes 1 μm. **(B)** Cumulative polypeptide binding over time (data extracted from the kymographs). Average values from 6 kymographs at 200 nM (green dots) and 5 kymographs at 50 nM (yellow dots) are plotted. The data is fitted with a Langmuir adsorption model resulting in a polypeptide binding constant of K ≈ 7·10^8^ M^−1^.

In a more detailed analysis of the kymograph data, we can determine that at 50 nM the peptides predominantly bind as trimers and at 200 nM as hexamers **(Fig. 5A)**. By analyzing the mobility of the polypeptides after binding, a strong correlation with cluster size is observed. Small clusters are able to slide along the DNA, while large clusters are effectively arrested on the DNA. By tracking single traces to obtain mean-square displacements (34) of polypeptide clusters moving on the DNA over time, we have obtained values of diffusion constants of polypeptide clusters bound to the DNA as a function of cluster size (**Fig. 5B)**. This quantitative analysis confirms that mobility of clusters drops to essentially zero for pentamers and higher order oligomers (*D* ≤ 0.1·10^−2^ μm^2^/s). The observed decrease of the diffusion constant with cluster size is much steeper than would have been expected for a simple linear scaling of the sliding friction with oligomer size (see **Fig. S2**). This suggests strong interactions with the DNA and possibly conformational integration of protein and DNA into growing VLPs.

**Fig 5.**
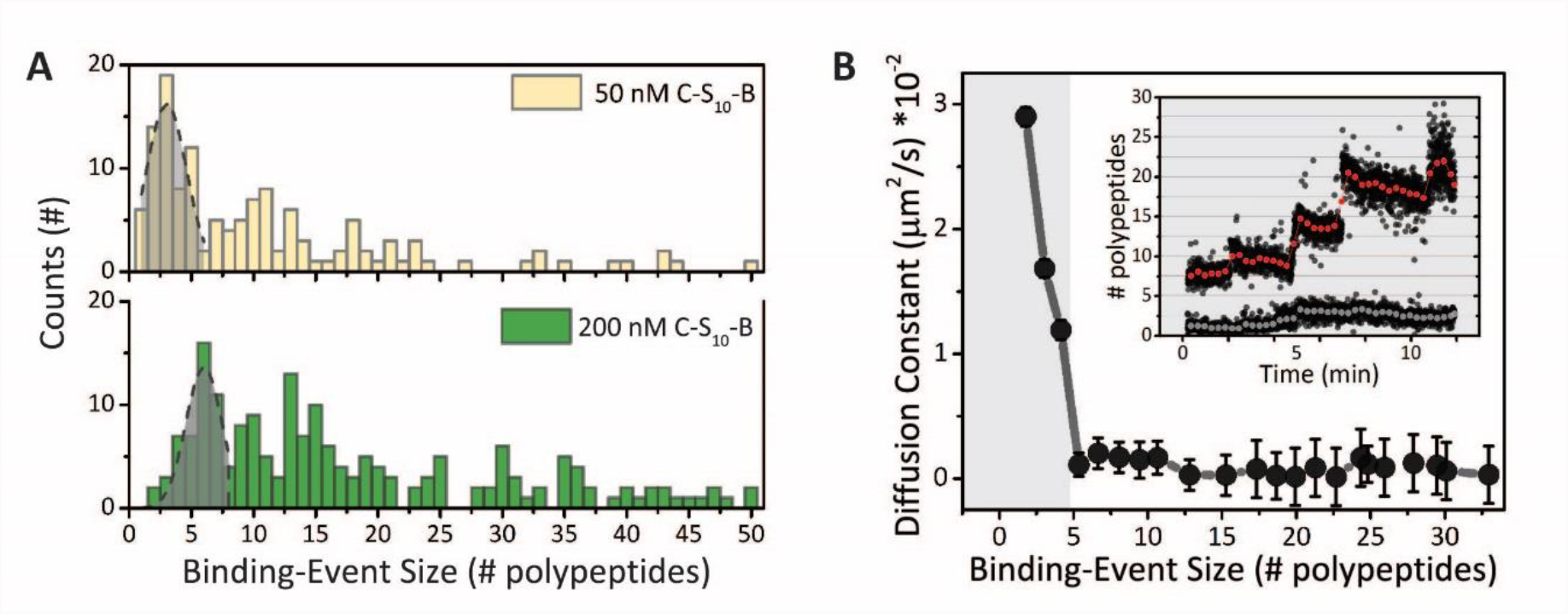
Polypeptide size quantification and their mobility along the DNA. **(A)** The quantification of single photobleaching steps allows to estimate the number of polypeptides bound per recorded event. The binding-event size statistics results in a histogram with a broad range of binding sizes. At 50 nM a majority of trimers is observed to bind (Gaussian peak, 3.0 ± 0.4 polypeptides). At 200 nM a majority of hexamers is found to bind (6.0 ± 0.3 polypeptides). **(B)** The diffusion constant *D* of the tracked binding events in the kymographs reveals an initial, drastic drop with increasing oligomer size, levelling off for oligomer sizes of ≥ 5 polypeptides (grey background area highlights the mobile events, error bars SEM). Inset: single binding events indicating that oligomer growth is more likely to take place when starting off with a large cluster (>5-mer, red curve) than with a small cluster (grey curve).

By quantifying the probability of growth (i.e., the increase in fluorescence intensity) of each event, we find that a polypeptide from solution is more likely to bind to a pre-bound polypeptide pentamer or bigger oligomer, with 90% of large clusters growing, while only 10% of the smaller ones increase in size **(Fig 5B, inset)**. The notion that a certain length of DNA needs to be covered by artificial capsid protein in order for VLP formation to commence is also supported by a bulk electrophoretic mobility shift assays (EMSA), which shows that binding of the artificial capsid polypeptides is strongly dependent on the length of the DNA template in the range of 10 bp to 1000 bp (**Fig. S3**).

Taken together, our data strongly suggests that pre-formed oligomers bind to the DNA, and that pentamers, when bound to the DNA, can be considered as critical nuclei for the productive formation of VLPs.

### Nature and dynamics of DNA condensation during capsid growth

The combined fluorescence microscopy and optical tweezers experiments on long DNA discussed above are not optimal for studying capsid growth. This is because i) the many nuclei on the long DNA (**Fig. S1**) make it difficult to follow growth of each of them individually, and ii) growth of the nuclei is a slow process that cannot be followed over sufficiently long periods of time due to photo bleaching. Therefore, as a complementary real-time, single particle technique, we use acoustic force spectroscopy (AFS). AFS allows us to probe end-to-end distances for short DNA molecules tethered between a surface and a microbead (see schematic in **Fig 6A**) as a function of time (up to hours) for a fixed low force and with high temporal resolution (50 Hz) (35). Since the DNA used in the AFS experiment is short (2.9 kbp ≈ 1μm), growth of virus-like particles is initiated from one or at most two nuclei only (**Fig. S1**), allowing us to follow growth in much greater detail. Furthermore, data for multiple DNA strands is acquired simultaneously, implying that high-throughput statistics can be obtained.

**Fig 6.**
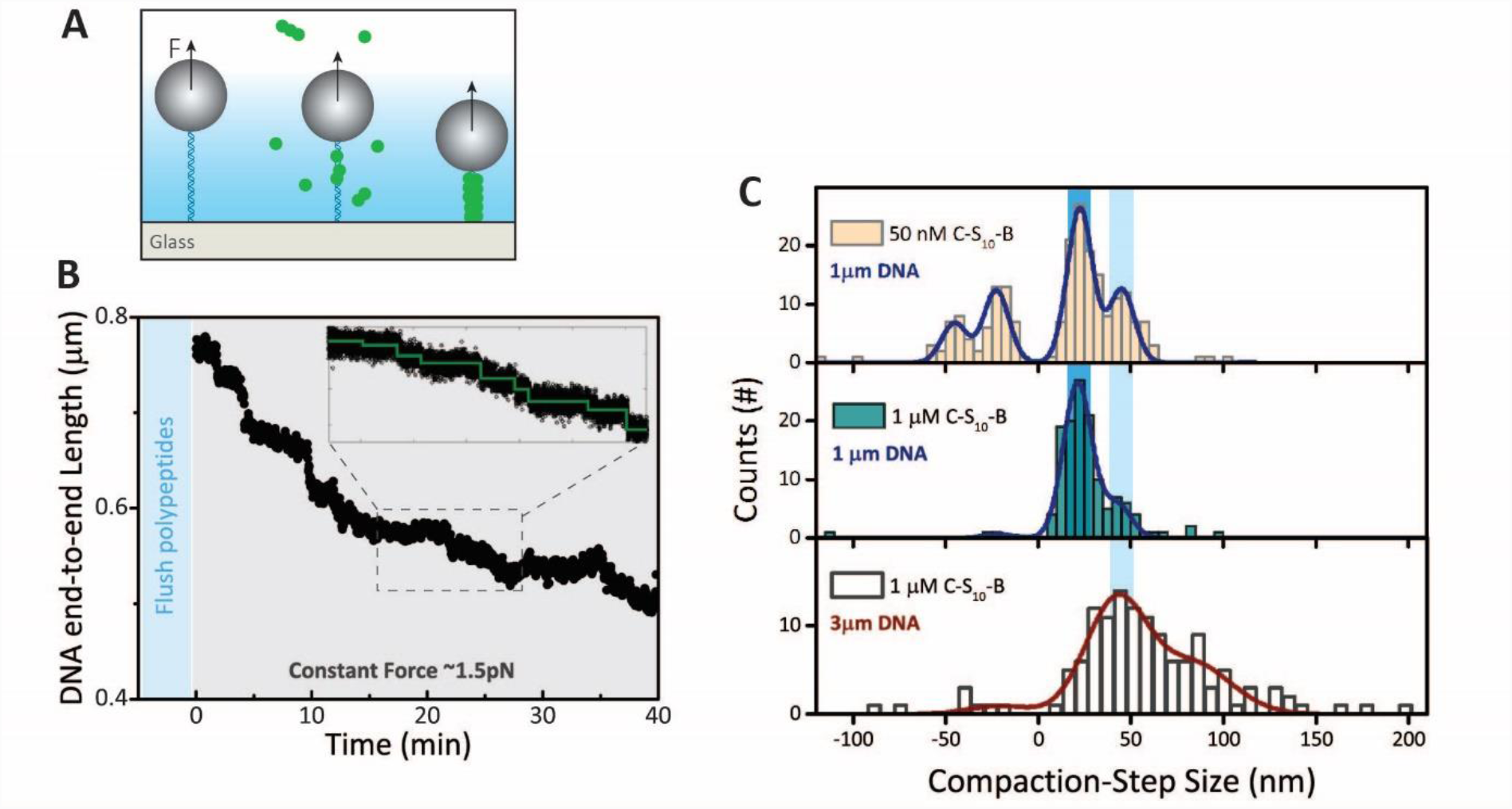
Acoustic force spectroscopy reveals particles compaction dynamics. **(A)** Illustrative image of the AFS setup. DNA tethered microspheres are pushed along an acoustically generated pressure gradient (blue/white background) allowing to apply a long and stable low-force clamp. **(B)** To monitor the particles compaction we record the DNA end-to-end length in the time. The light blue background indicated flushing in of peptides into the flow-cell while a stretching force of 15 pN was applied. The grey background area indicates a constant applied force of 1.5 pN. Inset: close-up of a compaction trace with the green line of the fit showing the compaction steps found with the step finding algorithm(36). **(C)** Step size statistics of compaction events at different conditions: 50 nM polypeptides-1μm DNA (top histogram), 1 μM polypeptides-1 μm DNA (middle histogram) and 1 μM polypeptides −3 μm DNA (bottom histogram). The negative steps obtained at lower concentration (top histogram) represent decompaction events which shows a symmetrical distribution. The compaction events data are fitted with a multi-Gaussian function, where the distances from peaks-to-peak are equally spaced and used as one fit parameter.

After confirming that results for effective persistence lengths deduced from force-extension curves obtained by OT agree with those obtained from force-extension curves obtained by AFS (**Fig. S4**), we start using AFS to probe in detail the nature and dynamics of DNA condensation during capsid growth. DNA condensation by the artificial capsid polypeptides is observed in real-time, by measuring the end-to-end distance of DNA molecules exposed to the artificial capsid proteins at a fixed low force of 1.5 pN. Surprisingly, we find that DNA condensation into virus-like particles proceeds in a step-wise fashion. We use a previously developed change-point analysis (36) to fit this data and extract step sizes **(Fig. 6B, inset**). To find most probable step sizes, we fit a multi-Gaussian distribution with equally spaced peak distances. The short DNA (≈ 1μm) reveals a sharp peak at a step size of 30 ± 1 nm in DNA contour length for both low (50 nM) as high (1 µM) C-S_10_-B concentrations **(Fig. 6C)**. This shows that the most probable step size for the condensation process is concentration-independent. At low polypeptide concentrations we also observe de-condensation steps, recorded as negative steps. Remarkably, the most probable step sizes for condensation and de-condensation appear to be equal, not only at low forces, but also when we increase the tension to induce de-condensation (**Fig. S4**). Employing a 3-fold longer DNA (8.3 kbp ≈ 3 μm) the step-size distribution has much less pronounced peaks, which we attribute to the presence of multiple growing nuclei on the longer DNA. In this case, simultaneous steps at multiple locations cannot be deconvoluted and are detected as larger steps **(Fig. 6C)**.

## Discussion

The self-assembly pathway of even relatively simple viruses, such as the tobacco mosaic virus that consists of a single-stranded RNA packaged by a large number of identical copies of coat protein, is highly complex. It is only partially understood and in fact remains the object of controversy (9). At least in part this is due to the fact that unresolved issues regarding capsid assembly pathways are difficult to address other than with real-time, single particle methods. Two such issues are considered in this work: the nature of the critical nuclei for productive capsid formation, and the dynamics and nature of nucleic acid condensation during capsid formation. For a simple artificial capsid polypeptide model system, which mimics essential features of the assembly of the much more complicated natural tobacco mosaic virus, we have shown that powerful real-time, single molecule techniques can be used to successfully address such issues.

For the artificial capsid polypeptides, we have used optical tweezers combined with confocal fluorescence microscopy to demonstrate that a broad range of pre-formed polypeptide oligomers can directly bind the DNA. As described by classical nucleation theory of protein capsids, a nucleus with a certain critical size has to be reached to trigger capsid formation (14, 37). We find that binding events of oligomers consisting of less than five polypeptides typically do not lead to particle growth. These oligomers, when bound, slide along the DNA with a mobility that rapidly decreases with increasing oligomer size. Binding events of oligomers consisting of minimally five polypeptides seem to be required for triggering particle growth. Such oligomers, when bound to the DNA are essentially immobile. Therefore, we conclude that the pentamers, bound to the DNA template, can be considered to be the critical nuclei for the formation of the artificial capsids. The smaller-sized oligomers (<5) that can slide along the DNA, may assists the growth process. Indeed, proteins sliding along a nucleic acid molecule during viral assembly are theoretically shown to accelerate considerably the self-assembly of natural icosahedral viruses (38, 39).

The nature and dynamics of DNA condensation during capsid growth was successfully addressed using AFS, since it allows probing end-to-end distances of multiple short DNAs over prolonged periods of time, under a precisely controlled low force. Surprisingly, we have established that DNA condensation into the artificial capsids occurs in discrete single compaction events, with approximately 30 nm of DNA contour length being condensed in each compaction event. This characteristic length of DNA per compaction event seems to be largely independent of the protein concentration. Also, de-condensation steps at low protein concentrations show the same characteristic length, suggesting that this length of DNA must corresponds to a characteristic structure of condensed DNA in the rod-shaped artificial viral capsid.

The filamentous core of the VLP is formed by the silk-like middle blocks S_10_ of the C-S_10_-B artificial capsid polypeptide, which assemble into a stack of beta-solenoids **(Fig. 7B)**. Each beta-solenoid sheet has a dimension of approximately 2.0 nm × 2.6 nm and a height of approximately 0.6 nm, as predicted by computer simulations (25, 40). The binding blocks B and stability block C emanate from the filamentous core. From this we expect that the DNA is confined to a condensation region extending at most a few nm away from the filamentous core **(Fig. 7)**, since the flexible oligolysine binding blocks B can only extend up to that distance. Such a structure is consistent with the height of the VLPs found using AFM imaging in liquid, which show an average particle height of ≈ 9 nm (**Fig. S1**).

**Fig 7.**
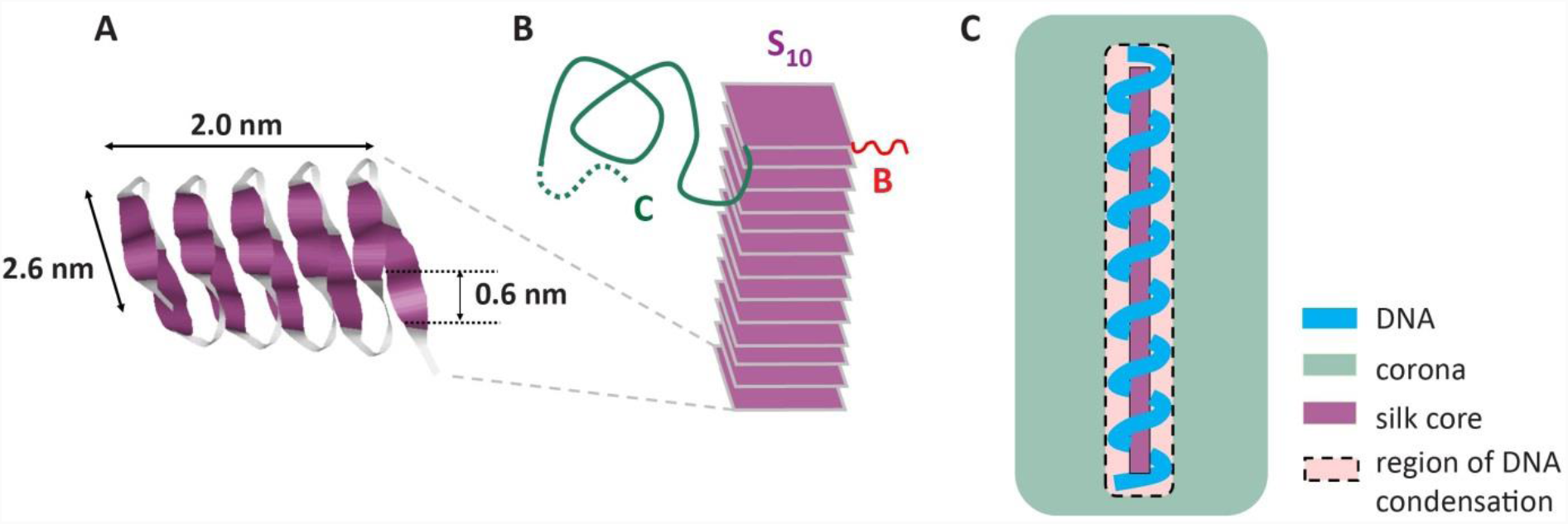
Conformation of condensed DNA. **(A)** Sheet-like beta-solenoid conformation of folded silk block S_10_ = (GAGAGAGQ)_10_ with approximate dimensions, as predicted by computer simulations (25). **(B)** The filamentous core of the VLPs is formed through stacking of the sheet-like folded silk blocks. **(C)** The region of DNA condensation extends from just outside the filamentous core up to the distance the flexible oligolysine binding blocks *B* = K_12_ can stretch away from the filamentous core from which they emanate, which is a few nm. Binding to the highly localized binding blocks may lead to different condensed conformations of the DNA such as a helical winding around the filamentous core of the VLP, as suggested by the observation of regular condensation and de-condensation steps of 30 nm of DNA contour length.

The question arises, however, what conformation the DNA adopts in the condensation region close to the filamentous core of the VLPs. For DNA packed into icosahedral spaces such as in T4 and T7 bacteriophages, it has convincingly been shown that the experimentally observed spool-like DNA configuration can be explained purely in terms of nano-geometric confinement (41–43). In other cases, binding to capsid proteins may induce nucleic acid template deformations that would not be expected on the basis of geometric confinement alone. For example, the helical arrangement of the RNA genome in TMV virus particles is dictated by their binding to the capsid proteins rather than by geometric confinement (44). For our artificial virus-like particles it has previously been shown that particle lengths are roughly one-third of the DNA contour length (21). If the DNA conformation inside the artificial virus-like particles considered here is determined by geometric nano-confinement alone, the most plausible conformation would be that of parallel double stranded DNAs with hairpin bending defects, as illustrated in **Fig. S5**. Such conformations minimize the bending energy of semi-flexible chains in finite length tubular confinement, for tube diameters much less than the persistence length, as predicted in recent computer simulations (45), and demonstrated by the theoretical estimates of **Eq. S10-S12**. Such conformations are also similar to the conformations adopted by DNA confined in nano-channels (46).

However, binding of the DNA to the highly localized binding blocks that emanate from the filamentous core may induce strong DNA deformations that may lead to DNA conformations very different from those predicted for confinement of the DNA in a finite nanotube. As was previously shown for several DNA-binding proteins (39,40), the effective persistence length reduction of the DNA we find upon the binding of the oligopeptide, suggests a deformation of the DNA backbone. Moreover, our finding of discrete DNA compaction events during capsid formation suggests that indeed for the VLPs considered in this work, deformations induced by the binding blocks, rather than a purely geometric confinement, determine the nature of the DNA condensation.

The observed 30 nm steps are suggestive of a stepwise helical winding of the DNA around the filamentous core of the virus like particle as it is growing during the AFS experiment, illustrated in **Fig. 7C** (see also **SI**). If we assume that the characteristic contour length of 30 nm corresponds to a single helical winding, this would imply a radius of the helix of 4.5 nm, and a radius of curvature of 5.1 nm, to arrive at a distance between the helical windings of 10 nm, consistent with the observed packing parameter of 3 (**Fig. S6**). This helical arrangement is large enough for the DNA to wind around the filamentous core of the VLP yet small enough to be within the region into which the binding blocks can extend. Interestingly, a DNA molecule wrapped around histones, has a similar radius of curvature (49).

An important caveat is of course that although our experiments are done under conditions quite close to assembly in free solution, the single-molecule methods that we use do impose some constraints that may influence the nature of the DNA condensation. In particular, we confine one of the ends of the DNA, and we always let assembly proceed in the presence of a small force. Nevertheless, the cooperativity of the C-S_10_-B artificial capsids assembly process (21) suggests that a regular growth from one location is the prevalent mechanism also when the compaction takes place with free DNA in solution, supporting the helical condensation hypothesis. Beyond the real-time single particle techniques that we have introduced here for studying the assembly of artificial and natural capsids, a next challenge would be to develop real-time microscopy of the growth of single capsids.

To summarize, for real-time, single particle level studies of the assembly pathways of artificial and natural viruses, the complementary techniques that we have used here are unique in combining high speed with single-particle level detail. We have used these techniques to address key issues regarding capsid assembly pathways that are difficult to address other than with real-time, single particle methods: the nature of the critical nuclei for productive capsid formation, and the dynamics and nature of nucleic acid condensation during capsid formation. Our study was performed on a simple artificial viral capsid protein but it paves the way for the detailed real-time *in vitro* studies of the assembly of natural viruses at the single-particle level.

## Materials and Methods

### Virus-like particle capsid polypeptides

The biosyntethic capsid polypeptides were provided as lyophilized protein polymer powder (C-S_10_-B = 44.7492 kDa), produced as previously described (21). We dissolve 0.5 mg/mL of the protein in demi-water, followed by 10 minutes incubation at 65°C. Next, we dilute the sample using a 10 mM phosphate buffer (pH 7.5) and supplement the sample with 0.1 mM dithiothreitol. For optical tweezers experiments C-S_10_-B was labelled with Alexa Fluor™ 555 C2 Maleimide (Thermo Fisher Scientific) that binds to the (only present) cysteine located at the N-terminal extremity of the shielding coil block C. The ‘Thiol-Reactive Probes’ protocol provided by the supplier was followed. The excess of unreacted dye was removed by columns filtration (PD-10 Columns G-25M). A Pierce BCA protein assay (Thermo Fisher Scientific kit) in combination with Nanodrop was then used for determining the protein and dye concentrations (ratio 1.5). The labelled protein compaction activity was tested both with AFM imaging and optical tweezers force measurements.

### Atomic force microscopy

Virus-like particles were imaged in Peak Force Tapping mode on a Bruker Bioscope catalyst setup, unless otherwise stated. Peptides and DNA were incubated with a final charge ratio N/P=3 (molar ratio between positively charged NH_2_ groups from the binding block, to negatively charged PO_3_ groups of the DNA template (P)), in 10 mM phosphate buffer at pH 7.5, for different timings (see main text). Virus-like particles were adhered to freshly cleaved mica treated with a 5 mM TRIS and 0.5 mM Mg^+2^ solution. For the AFM imaging in air, the silica substrate was cleaned with 70% ethanol followed by acetone and dried under nitrogen flow. Then the sample was deposited onto the surface for 2 minutes, afterwards the slides were cleaned with MQ-water and dried slowly under N_2_ stream. Liquid AFM experiments were performed in 10 mM phosphate buffer at pH 7.5 on freshly cleaved mica. Silicon nitride cantilevers (Olympus; OMCL-RC800PSA) were used, with a nominal tip radius of 15 nm and spring constant 0.76 N/m and 0.10 N/m for air and liquid experiments, respectively. Individual cantilevers were calibrated using thermal tuning. AFM image processing was performed with NanoScope Analysis 1.5 software for both a first order imaging flattening and the particles height estimation. For nucleation site quantification and for the images in Fig. S1A a Digital Instruments multimode AFM with a NanoScope V controller and silicon nitride cantilevers with a spring constant of 0.4 −0.35 N/m (SCANASYST-AIR, Bruker, MA, USA) was used. Protein-DNA samples ([2.5 kbp dsDNA] = 1 µg ml^−1^ or 0.65 nM, and [Protein] = 121.6 µg ml^−1^ or 2.692 µM) at N/P = 10 in 10 mM phosphate buffer, pH 7.4 with 0.1 mM DTT, were deposited and incubated during 2-3 minutes onto a clean silicon surface, previously treated in a plasma cleaner for around 2 minutes. Immediately, the substrate is rinsed with 1 mL of Milli-Q water, and then excess of water is removed by soaking up using a tissue and slow drying under a N_2_ stream. To process the images we used NanoScope Analysis 1.20 software. A first order flattening was applied to all images. We perform contour length measurements using ImageJ software to extract the number and length of the nucleation points on the DNA single molecules.

### Optical tweezers with confocal fluorescence microscopy

The dual-trap optical tweezers set-up with integrated confocal fluorescence microscopy (LUMICKS), is similar to an optical set-up used for dual-trap optical trapping experiments in combination with confocal fluorescence has been described previously (50). Microfluidics: a 5-channel laminar flow cell (LUMICKS) was assembled onto an automated XY-stage (MS-2000, Applied Scientific Instrumentation). The latter allowed the controlled transfer, in a highly efficient manner, of the optically-trapped beads through the channels of the flow cell, containing the molecules of interest. End-biotinylated bacteriophage *λ* DNA was connected to streptavidin-coated polystyrene beads (Ø = 4.5 μm, Spherotech) to generate the DNA constructs, as described previously(51). Protein–DNA interactions and force-stretching experiments were conducted in 10 mM Phosphate buffer (pH 7.5) and 0.1 mM dithiothreitol. Kymographs were recorded by scanning the confocal spot along the DNA kept at a fixed distance (corresponding to 5 pN on bare DNA), with a scanning-line time of 200 ms (averaged over three lines during analysis and resulting in a temporal resolution of 600 ms). Binding of single peptides was followed through kymographs analysis quantifying their fluorescence signal (average number of photons) when landing on the DNA. All values were background corrected. We processed the kymographs through single-molecule tracking to acquire information on the binding events intensity and mobility. Photobleaching allows to calibrate the intensity of a single fluorophore (11.1±0.5 photons) by looking at single fluorescence decrease steps of single photobleached dyes. The one-dimensional diffusion of protein complexes along the DNA was quantified by tracking the peptides traces for a large number of scanning-lines (more or equal to 300) and calculating their diffusion coefficient (*D*) by using a mean square displacement analysis (MSD) (34). Force-distance curves and confocal fluorescence data were analyzed using a custom-written MATLAB software. Here the extensible worm-like chain model (eWLC) (52), which describes the dsDNA elastic behavior up to ∼30 pN, is used to fit FDCs and estimate the DNA effective persistence length 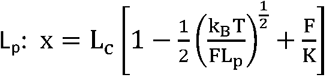.

### Acoustic force spectroscopy

The home-built AFS setup (35, 53) and the AFS flow-cell (LUMICKS) and tethers preparation (53) have been previously described. The 8.4 kbp DNA was obtained from a pKYBI vector, as previously described(53).. The shorter dsDNA is obtained from a 2.9 kbp pRSET-A plasmid (Thermo Fisher Scientific, V35120) was fabricated in the same way as the PKYBI construct except for the last purification step that is not performed. About 1 μg (7μL of a concentration of 149 ng/μL) of plasmid is used for the first preparation step. pRSET DNA has KpnI and EcorRI cutting sites as well. Following the same steps, a volume of 50 μL with a final concentration of 6 ng/μL is obtained. This construct was pre-incubated with the microspheres: ∼50 pg of pRSET-A DNA was mixed and incubated for 10 min with 10 µL of 4.5 µm streptavidin coated polystyrene beads. The microspheres were cleaned by adding 1 mL of PBS (138 mM NaCl, 2.7 mM KCl and 10 mM phosphate (pH 7.4)) supplemented with EDTA (5 mM), casein (0.02% w/v), pluronics F127 (0.02% w/v) and Na-azide (0.6 mg/mL), called measuring buffer. Next, the sample was spun down (2000 g for 2 minutes) and at last the residue was removed. This step was repeated three times and the final product was supplemented with 20 µL with measuring buffer. This was flushed in the AFS and after 30 minutes of incubation, the free microspheres were flushed out with measuring buffer and the construct is ready for measuring. AFS data were analyzed using a custom-written LABVIEW software, and the step-analysis was performed with a custom-made change-point analysis software (36). Processed data were analyzed using Origin. Gaussian fit in Fig. 6C: The peak-to-peak distance obtained are 22.6 ± 2, 21.3 ± 0.4 and 21 ± 0.3 nm for the histogram from top to bottom. The light blue backgrounds highlight the mean of the first two Gaussian peaks. In the main text these values are corrected for the force applied during the experiments and the observed change in the effective L_p_, resulting in an average step of 30 nm.

### Electrophoretic Mobility Shift Assay (EMSA)

EMSAs were performed to determine the effect of the dsDNA length on the protein binding. Samples were prepared in the following way: water protein C_4_S_0_-_10_B solution, previously heated to 343 K for 10 minutes, was added to a solution containing 4 ng/μL dsDNA (10, 100, 1000 or 2000 bp) in 10 mM Tris buffer at pH 7.4 and incubated during 60 minutes at room temperature. Then the samples were loaded on 20% acrylamide gels in 1x TAE buffer and run at 70V for 90 min. Gels were in a gel documentation system and analyzed with ImageJ. N/P ratio for 50% binding of DNA (K_Dapp_) by the protein was calculated fitting the DNA free intensities to the Hill equation, fraction bound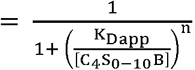, with n the Hill constant.

## ACKNOWLEDGMENTS

This work was supported by a STW HTSM grant (to GJLW and WHR) and a NWO Vidi grant (to WHR). We thank Daan Vorselen for providing the steps-finding algorithm, Denise Denning for performing AFM measurements in liquid and Andreas Biebricher for support with AFM experiments.

## Supplementary Information

### Atomic force microscopy imaging

**Fig S1.**
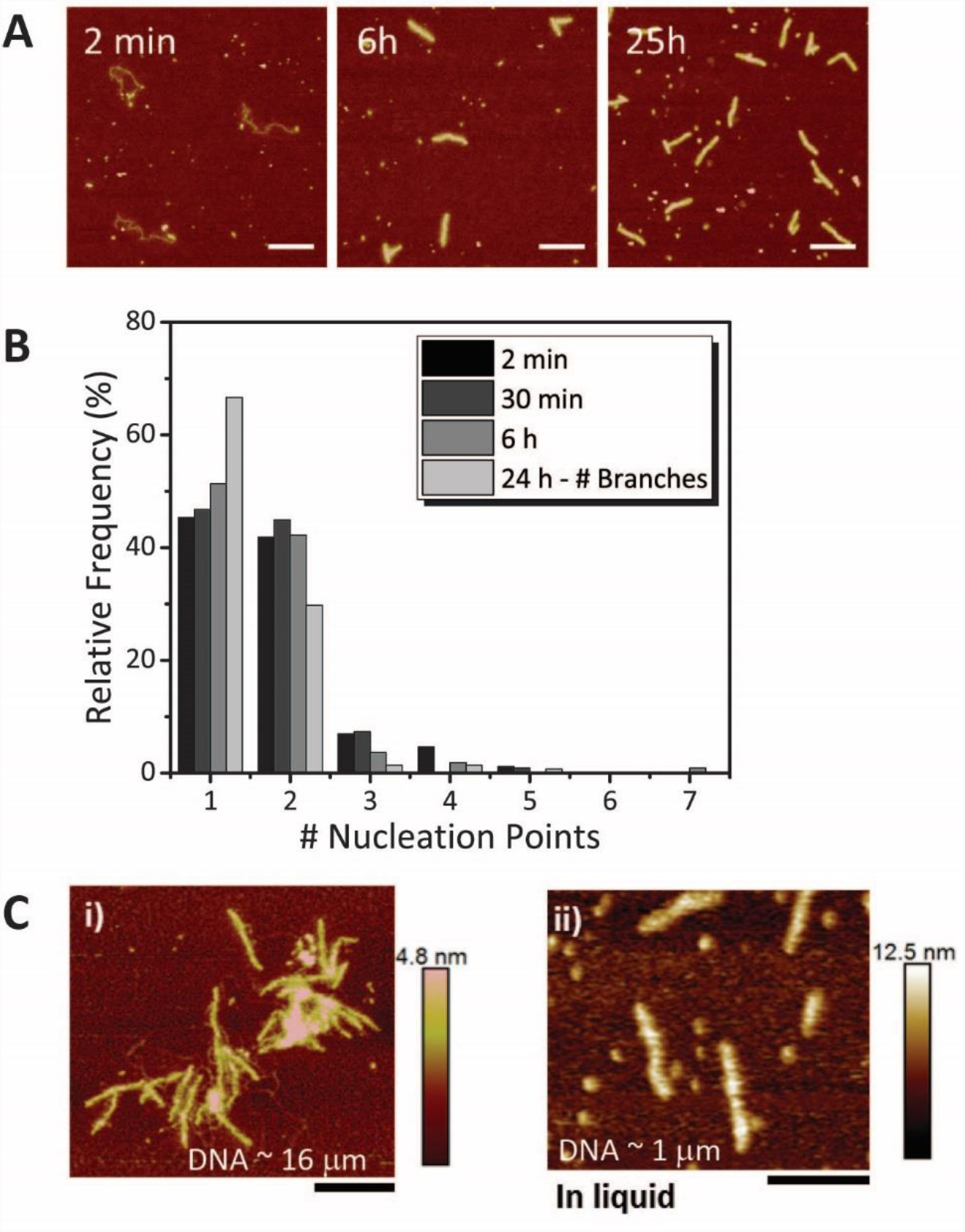
**(A)** AFM images taken at different incubation times of capsid polypeptides with 2.5 kbp-long DNA molecules (see Methods). Scale bar 350 nm. **(B)** Quantification of the number of observed nucleation points along single DNA molecules at different incubation times (N= 86, 109, 109, 141 for 2 min, 30 min, 6 h and 24 h, respectively). Note: after 24 h incubation a full particle is formed thereofe it is possible to visualize the particle branches instead of nucleation sites. **(C)** Particles on a 16 μm-long DNA, used for optical tweezers experiments and imaged in air. Multiple nucleation sites result in multiple particles formed on the same DNA (Panel i). Imaging in buffer solution results particles that are ∽4 times higher than in air. The average height in liquid is 9 nm (Panel ii). Scale bar 300 nm.

### Optical Tweezers with confocal fluorescence experiments

**Fig S2.**
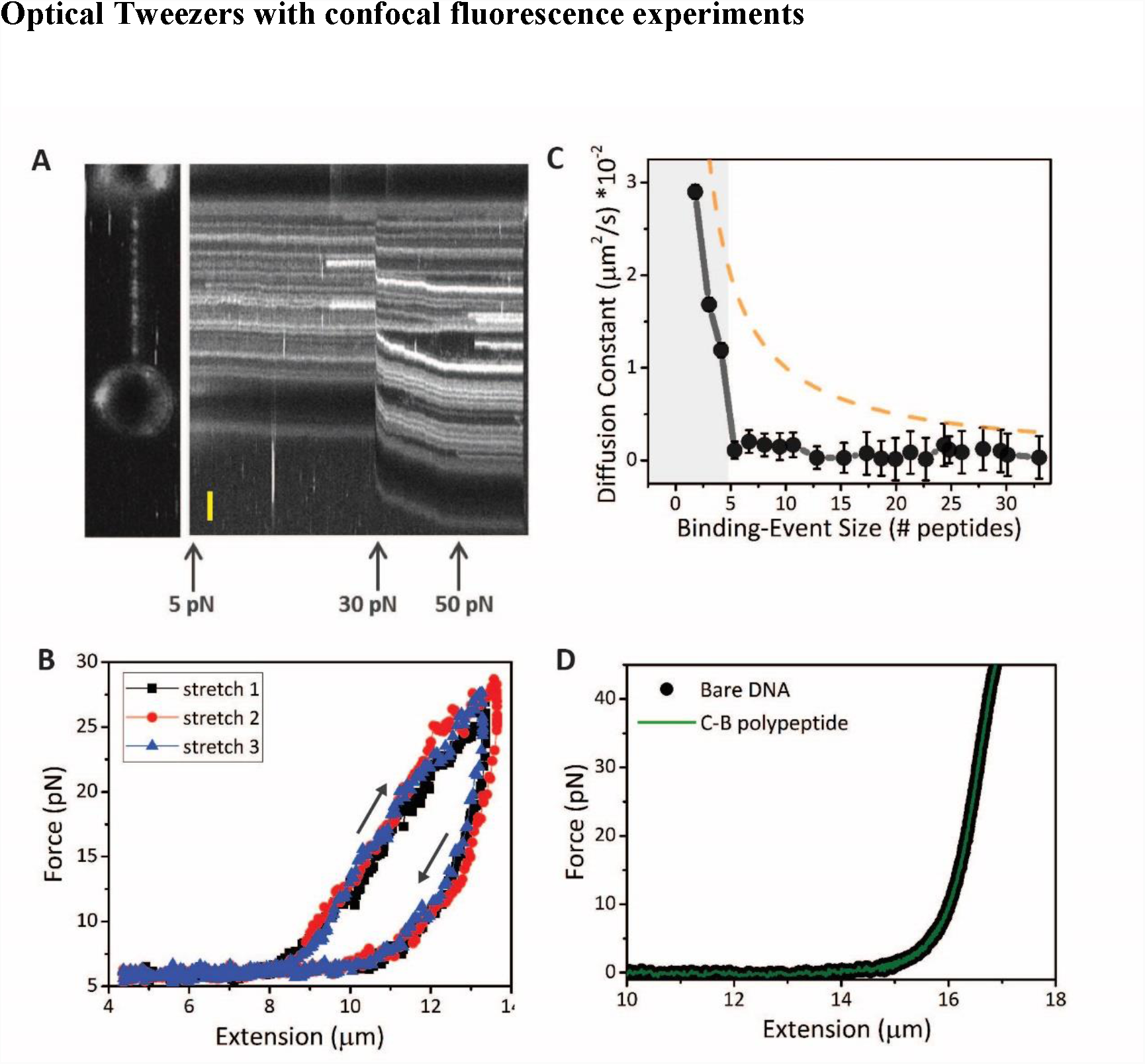
(A) Confocal fluorescence image and corresponding kymograph of a pulling experiments at high force (> 40pN), which shows that peptide unbinding is not observed, and peptides binding can continue to take place at high DNA tension. (B) Repetitive stretch-retraction curves of DNA coated with C-S_10_-B, in the absence of polypeptides in solution. The plot shows that the particle compaction is restored when going back to low forces (i.e. overlapping between three repetitive curves). (**C**) Diffusion constant as function of the binding event size (black data identical to Fig. 5B in main text) and comparison with expected diffusion constant decay if only the mass increase would affect the change in diffusion (dashed, orange line). This theoretical decay does not follow the experimental data, thus revealing that the observed drastic drop in the diffusion constant cannot be fully explained by the mass increase. Together with other evidences (see main text) suggests that a nucleus is formed when more than 5 units bind the DNA. **(D)** Force distance curves of bare DNA (black) and DNA incubated with C-B polypeptide (green) lacking the central silk-block. The DNA mechanical response is unaffected in the presence of C-B which does not compact the DNA neither contribute to a change in its effective persistence length.

### Bulk assay

**Fig S3.**
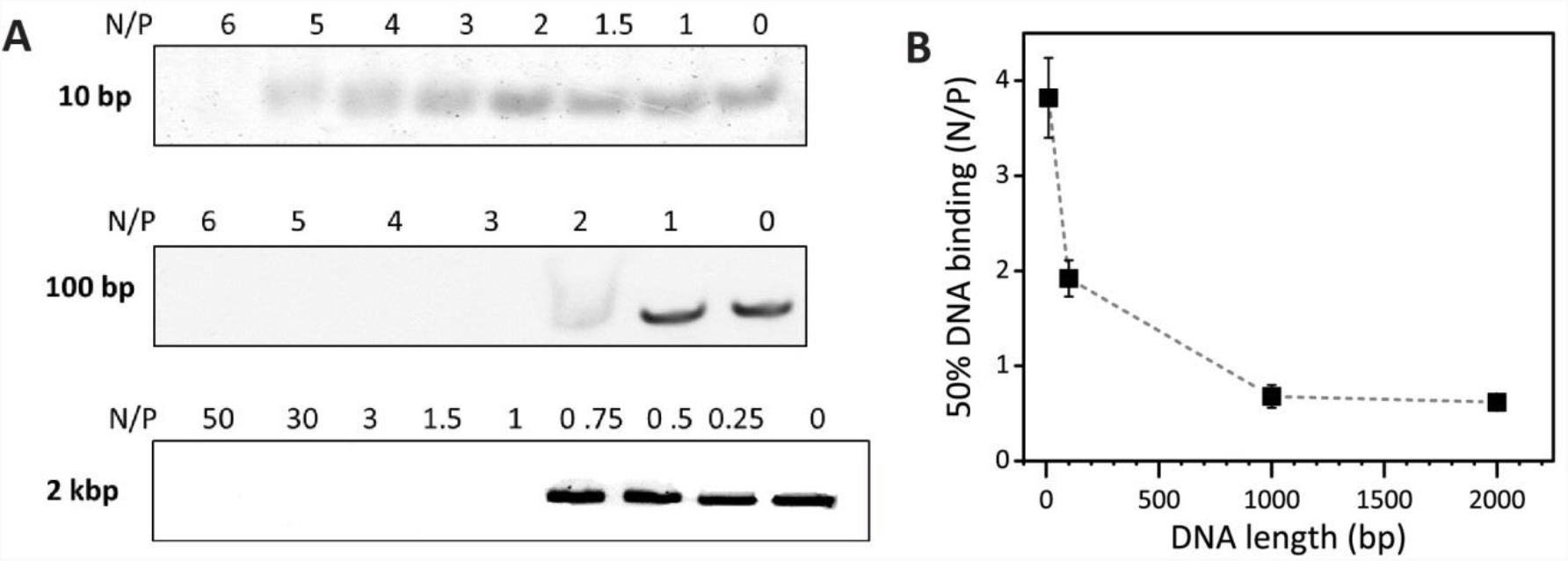
**(A)** Gels showing electrophoretic Mobility Shift Assay (EMSA) of DNA with length of 10, 100 and 2000 bp when incubated with protein C-S_10_-B. **(B)** Protein concentration (as N/P) to achieve 50% DNA binding (related to protein affinity) is plotted over function of DNA length.

### Acoustic force spectroscopy experiments

**Fig S4.**
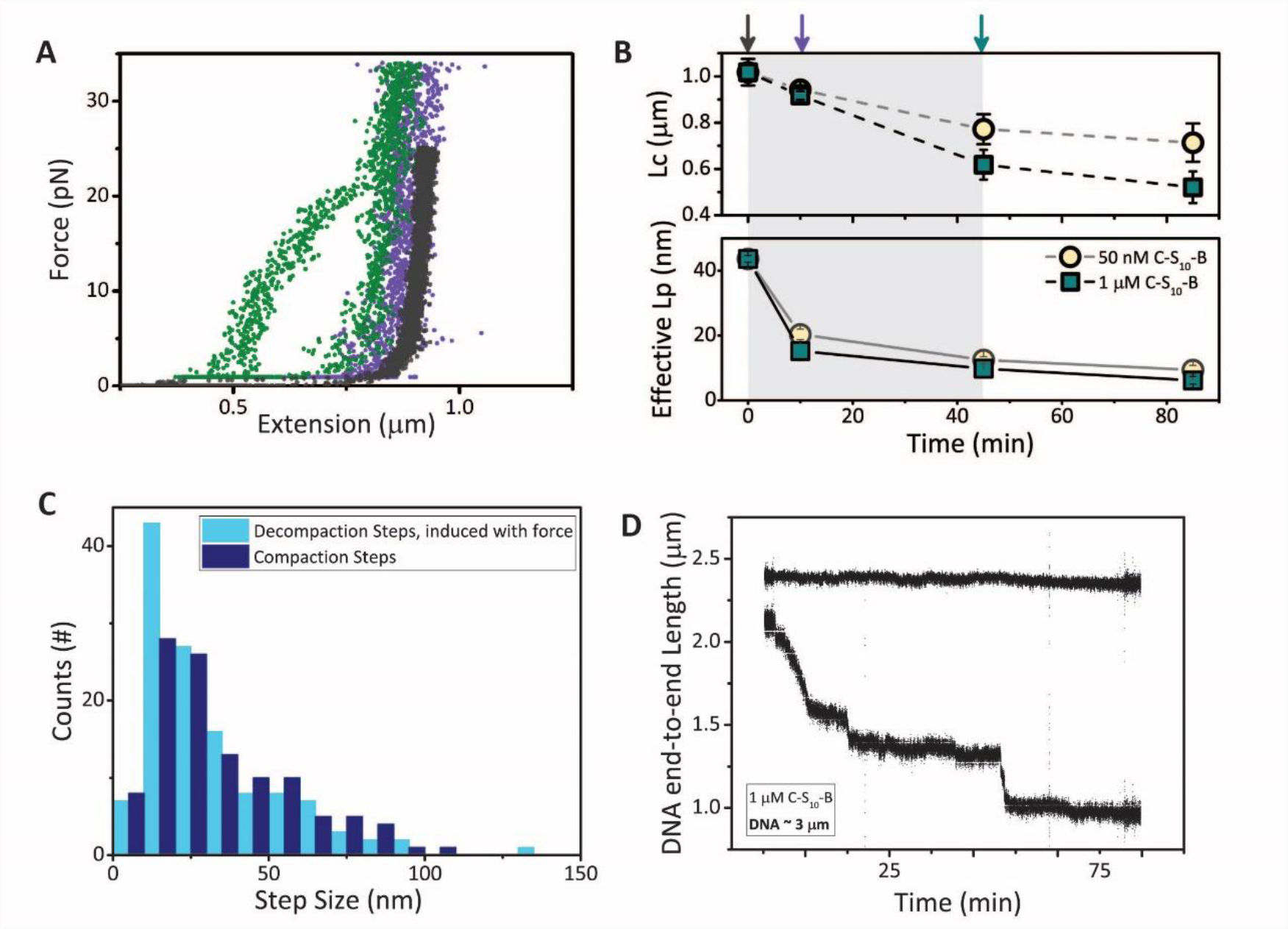
AFS additional observations. **(A)** Representative AFS force-distance curves. **(B)** The contour (top) and effective persistence (bottom) lengths are obtained from fitting WLC to the forward-and the back-stretching curves, respectively. A total of 20, 20, 15 and 9 curves were fitted for 50 nM polypeptide concentration and 15, 15, 15, 12 and 8 for 1 µM concentration, for curves obtained after 0, 10, 45 and 85 minutes, respectively (error bars are SEM). The results are in agreement with optical tweezers force experiments. **(C)** Force-induced particles decompaction (dark blue histogram) shows the same steps sizes as shown during the compaction events (cyan histogram), constant force used 20 pN. **(D)** Two examples of DNA end-to-end length recorded during the compaction of ∽3mm-long DNA at 1mM capsid polypeptides concentration. While the top molecule is not being compacted the bottom one continues growing; this indicates that, as previously observed by Hernandez et al. (1), the artificial virus-like particles follows a cooperative assembly.

### Conformation of condensed DNA in artificial viral capsids

If geometric nano-confinement alone governs DNA condensation in the VLPs, one would expect condensed conformations of parallel double stranded DNA and hairpin bending defects, as illustrated in **Fig. 7D**. This has been found in computer simulations for semi-flexible polymers confined in narrow tubes (2) and is also expected on the basis of scaling arguments: parallel packing with a threefold compaction in length (or in other words, with a packing factor *p* = 3) as expected for the VLPs in principle requires no more than two hairpin bending defects. If the radius of the region of DNA condensation is *r*, the energy ϵ_*loop*_of such hairpin bending defects is

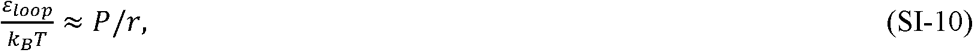

which is at most *O*(10*k*_*B*_*T*), where *k*_*B*_*T* is the thermal energy. In contrast, if we assume a helical packing as in **Fig. 7E**, for a helix of radius *r* the total bending energy is extensive in the length *L* of the template,

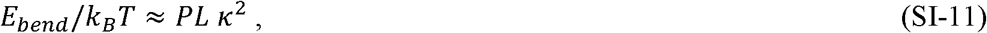

where κis the curvature of the helix,

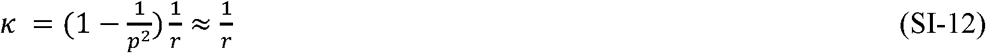

and *p* = 3 is again the packing factor of the VLPs (which equals the ratio of the arc length over the height of the helix). For large enough template lengths *L* the non-extensive bending energy of hairpin defects will always be smaller than any extensive bending energy associated with uniform bending as in a helical conformation. Indeed, for typical numbers *L* = 300 nm, *P* = 50 nm, *r* = 5nm, we arrive at bending energies *O*(100 *k*_*B*_*T*). Hence, purely geometric nano-confinement would not favor a helical arrangement of condensed DNA, but the bending energies of helical packing can be easily offset by the lowering of the free energy due to the binding of DNA to the oligolysine binding blocks *B*, similar to the binding to-, and induced wrapping of DNA around histones.

**Fig S5.**
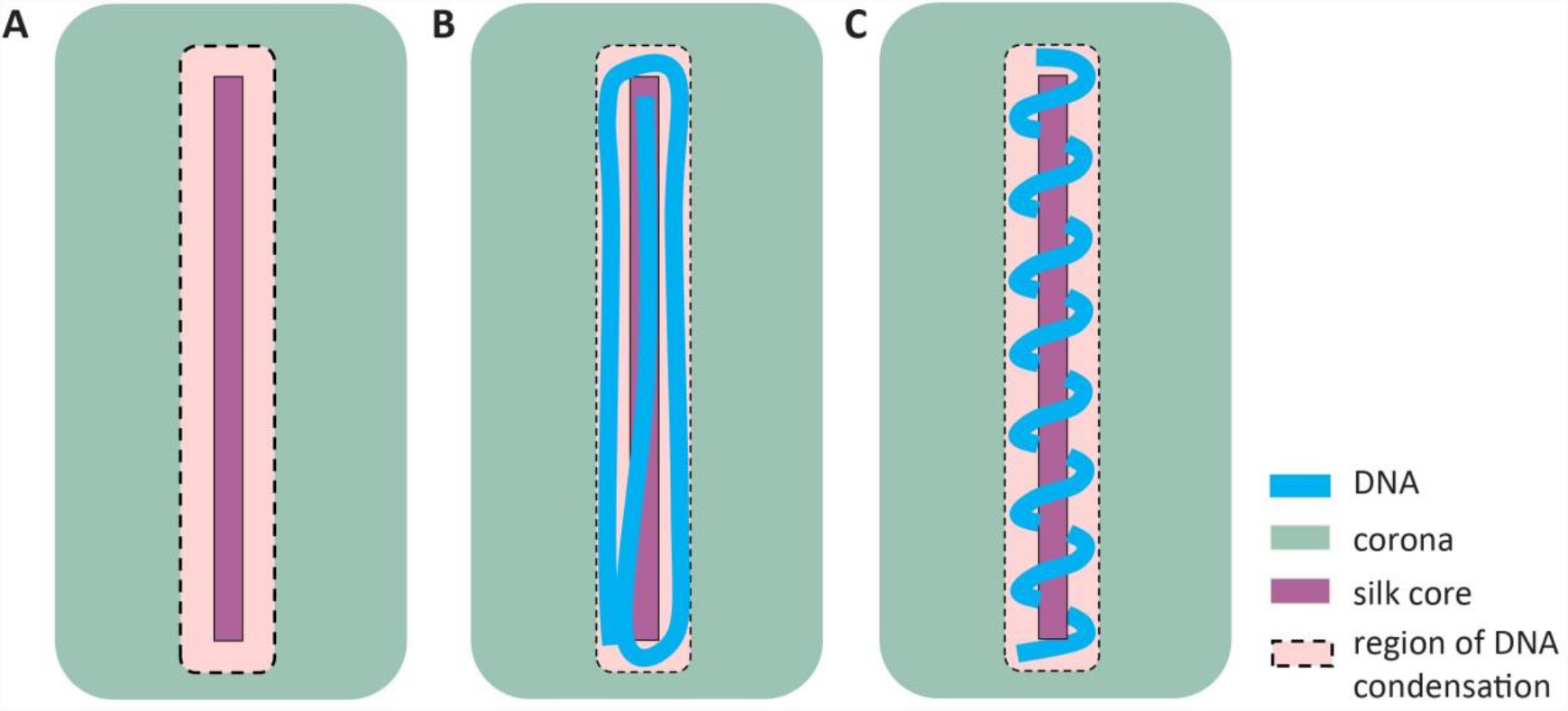
**(A)** The region of DNA condensation extends from just outside the filamentous core up to the distance the flexible oligolysine binding blocks B = K12 can stretch away from the filamentous core from which they emanate, which is a few nm. **(B)** If condensation is governed by geometric nano-confinement into a narrow tubular space, the expected conformation would be that of parallel double stranded DNA’s with hairpin bending defects. **(C)** Binding to the highly localized binding blocks may lead to different condensed conformations of the DNA such as a helical winding around the filamentous core of the VLP, as suggested by the observation of regular condensation and decondensation steps of 30 nm of DNA contour length.

**Fig S6.**
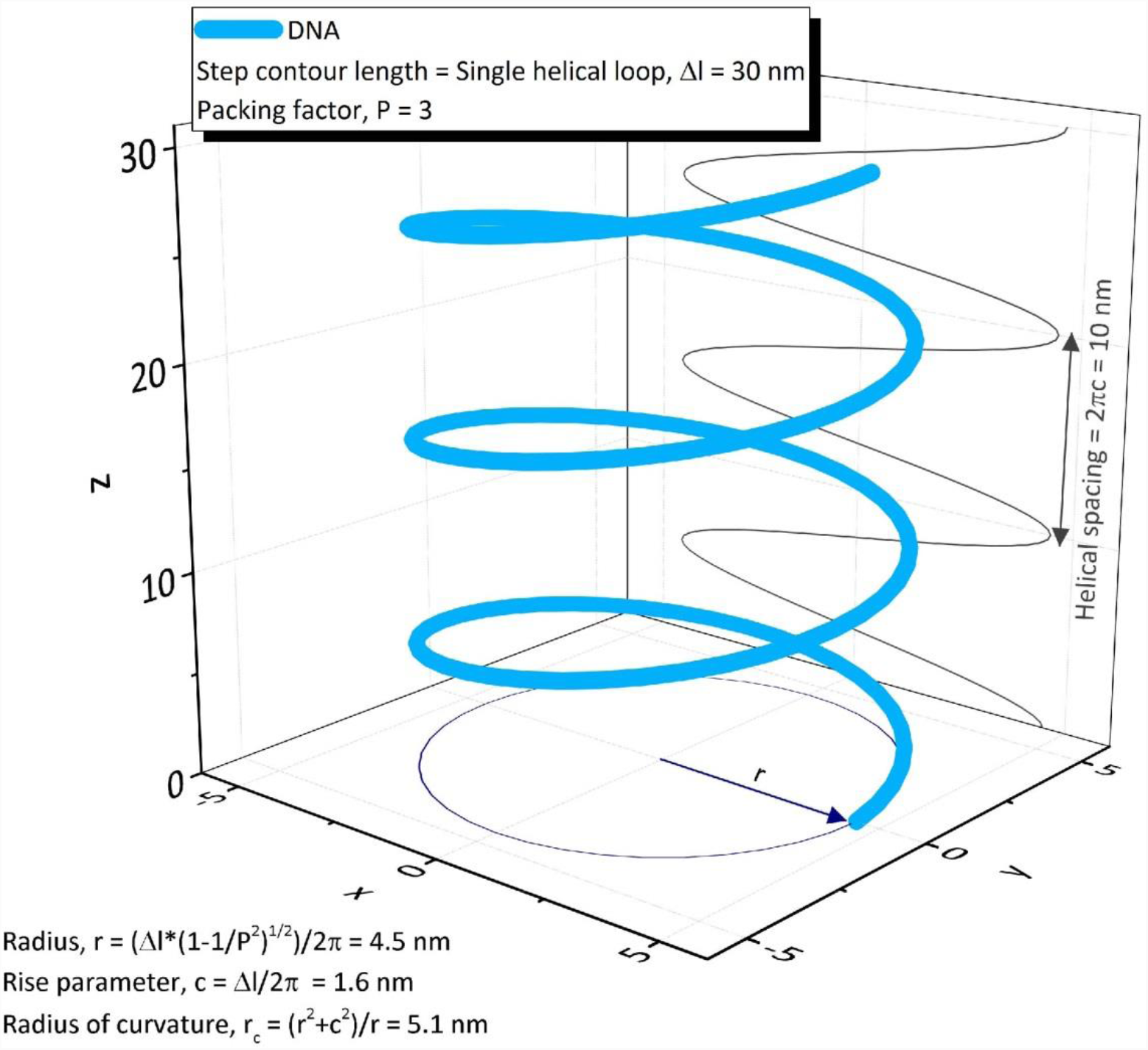
Helix representation of a DNA (light blue) with contour length of 90 nm, which is compacted in helical loops of 30 nm (i.e. the observed compaction step size), to a final packing factor of 3. The helical parameters described in the main text, are derived from the equations in the plot.

### Langmuir dynamics

The Langmuir adsorption equation is obtained as follows: Consider a surface, e.g., of a DNA molecule, in contact with a reservoir containing a mole fraction *X* of a solute, e.g., a protein, in a dilute (ideal) solution. The reservoir is so large that any binding to the surface does not appreciably deplete the solute from it, in other words, its concentration remains an invariant of time. Let *θ*(*t*)be the fraction of binding sites occupied by a protein at time t. If every binding site can bind a single solute particle, and the solute particles bound to the surface do not interact with each other, it seems reasonable to presume that Langmuir dynamics applies. In other words, *θ*(*t*)obeys the following kinetic equation

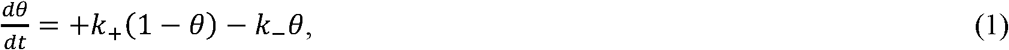

with *K*_+_the forward (binding) rate and *K*_-_the backward (unbinding) rate. The forward rate *K*_+_does not depend on the mole fraction *X* of solute in the solution, if binding is reaction-limited and involves a free energy barrier, e.g., due to a necessary conformational adjustment before binding is possible. If this is not the case, then the forward rate must be diffusion limited, and we expect *K*_+_∝ *X*. The backward rate *K*_-_presumably does not depend on the concentration of solute in the solution, and if the bound particles do not interact then it should be an invariant of *θ*(t).

If the steady-state binding and unbinding approaches a state thermodynamic equilibrium, then *K*_+_and *K*_-_are not independent. Indeed, under conditions of steady state, 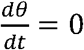, we have from eq (1)

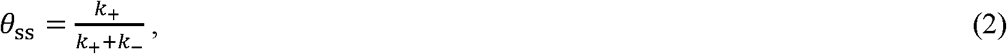

implying that if there is no unbinding, in other words *K*_-_= 0, that *θ*_ss_= 1 and all sites will become occupied. If the steady state is a state of equilibrium, *θ*_ss_should be equal to the equilibrium bound fraction *θ*_eq_, which obeys Boltzmann statistics,

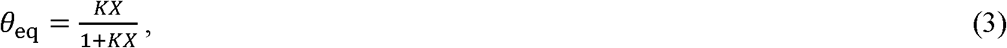

where *K* is the (dimensionless) binding constant. Eq (3) is the well-known Langmuir isotherm. Hence, under conditions of reversible adsorption, we must have

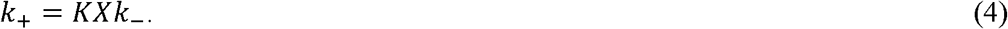

Eq (1) can be solved exactly, e.g., by variation of constants, to give

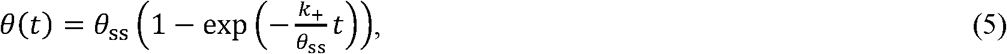

implying that the characteristic adsorption timescale is given by τ = *θ*_ss_/*K*_+_. For reversible adsorption, this can be written as

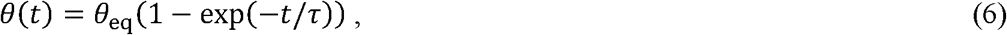

where relevant timescale becomes τ = *θ*_eq_/*K*_+_.

It is useful to consider two regimes. First, take conditions where *KX* ≫ 1, so concentrations much larger than the “critical” concentration 1/*K* needed to occupy half of the total number of binding sites. In that case, *θ*_eq_∽ 1 – 1/*KX*→ 1 and

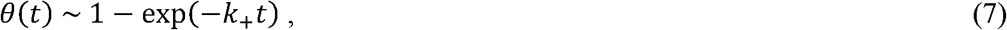

leading to full coverage at long times t ≫ 1/*K*_+_. Note that in this limit, the saturation value of the adsorbed amount is very weakly dependent on the concentration of solute.

For concentrations much below the critical concentration we have *θ*_eq_→ *KX*, implying

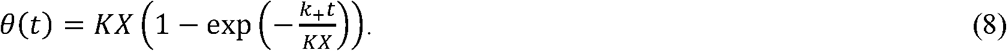

Equation implies the existence of three rather than two parameters, because *θ* = *N*/*N*_max_with *N* = *N*(*t*)the number of bound molecules (per unit area) and *N*_max_the maximum number of bound molecules (per unit area). The latter is equal to the number of effective binding sites. Note that for short times, that is, for t ≪ *KX*/*K*_+_, we can rewrite eq (8) into

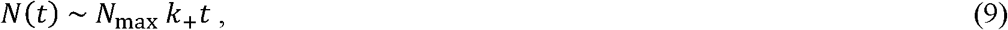

suggesting that, if data taken at sufficient numbers of concentration, curve fitting should allow us to fix the values not only of *K*_+_and *K*, but also that of *N*_max_.

1. Hernandez-Garcia A, et al. (2014) Design and self-assembly of simple coat proteins for artificial viruses. Nat Nanotechnol 9(9):698–702
2. Fritsche M, Heermann DW (2011) Confinement driven spatial organization of semiflexible ring polymers: Implications for biopolymer packaging. Soft Matter 7(15):6906.

